# Label-free visualization of ciliary rootlets in mouse brain

**DOI:** 10.1101/2024.06.19.597702

**Authors:** Yusuke Murakami, Mutsuo Nuriya, Zuliang Hu, Masaki Tomioka, Ryosuke Oketani, Kotaro Hiramatsu, Philippe Leproux, Akihito Inoko, Sakiko Honjoh, Hideaki Kano

## Abstract

Neuronal primary cilia are important role in brain development, sensory perception and neurogenesis. Rootletin, a fibrous protein composed of coiled-coil motifs, is a major structural component of ciliary rootlets and is essential for understanding ciliary functions. However, the precise mechanisms by which Rootletin influences ciliary dynamics and impacts neuronal function remain largely unknown, primarily due to the challenges in visualizing these structures. Here, we describe a label-free, rapid, and highly sensitive method to visualize Rootletin molecules in brain tissue. This platform integrates a second harmonic generation (SHG) microscope and background reduction by a one-step chemical pretreatment. Additionally, we employ coherent anti-Stokes Raman scattering imaging to simultaneously determine the cellular regions and intracellular locations of SHG signals. By applying this multimodal multiphoton imaging to mouse hippocampus, we found that neuronal ciliary rootlets were found to exhibit highly organized specific intracellular distributions. Moreover, the formation of ciliary rootlets precedes that of primary cilia. These findings highlight the utility of our label-free imaging platform in developmental and neuroscience research, providing a new tool to characterize ciliary dynamics and neuronal function.

## Introduction

Primary cilia play a key role in recognizing the physical and chemical parameters outside the cell and consequently maintaining cellular homeostasis (1, 2). Neuronal primary cilia, in particular, are essential in signal processing (3, 4), including cell polarity (5), axonal guidance (3, 6, 7), and neurogenesis in the brain (8-10). Dysfunction of neuronal primary cilia can cause numerous adverse outcomes, including neurodevelopmental disorders (11, 12), cognitive impairment (13, 14), and schizophrenia (15, 16). These ciliopathies are at least partially attributable to structural abnormalities in the ciliary rootlets, which extend from the basal bodies and anchor the cilia to the cell membrane (17, 18). Accordingly, understanding the molecular basis of abnormal neuronal primary cilia formation has become a research focus, with significant efforts dedicated to studying Rootletin, the primary structural component of ciliary rootlets. For example, studies using Rootletin knock-out and knock-down cells have demonstrated a significant decrease in the cilia formation rate (19), leading to reduced primary cilia formation (20).

Despite extensive research on the relationship between neuronal primary cilia, ciliopathies, and various brain functions, the molecular mechanisms by which Rootletin influences ciliary dynamics and impacts neuronal function remain largely unknown. This is partly due to the lack of reliable methods for visualizing ciliary rootlets with sufficient spatial resolution and reliability. Traditionally, ciliary rootlets can be visualized by immunostaining of Rootletin (18-20). How ever, this method is limited by nonspecific binding and poor reproducibility due to complicated interactions between the antibodies and the target proteins. Furthermore, for diagnostic purposes, the time-consuming immunostaining process hinders its broad application in clinical practice.

In this study, we demonstrate the selective second harmonic generation (SHG) imaging of ciliary rootlets by integrating our custom-built, highly sensitive SHG microscope with a background reduction technique comprising a one-step pretreatment. Our spectrum-based SHG microscopy enables the visualization of non-centrosymmetric structures in small objects (21), facilitating label-free detection of ciliary rootlets (22). Meanwhile, major brain components, such as axons (23, 24), also exhibit strong SHG signals, contributing to background signals. Hence, this background signal is suppressed using a one-step paraformaldehyde (PFA) treatment, thereby achieving the selective visualization of ciliary rootlets. Additionally, we simultaneously acquired coherent anti-Stokes Raman scattering (CARS) images, providing information regarding the cell regions and intracellular localization of SHG spots. By analyzing the positions of the ciliary rootlet relative to the cell body, we found that neuronal ciliary rootlets exhibited highly organized intracellular distributions. We further applied our method to the characterization of neuronal development by acquiring SHG/fluorescence images at different timings after the differentiation, where the results indicate that the formation of ciliary rootlets precedes that of primary cilia by at least few days. The present method can be applied characterize brain formation in a multi-scale context from organelles to tissues.

## Results

### SHG provides label-free ciliary rootlet imaging through PFA treatment

We first demonstrated SHG-CARS imaging (Figure 1) of selective ciliary rootlet in mouse hippocampus densely covered with axons. To achieve this, we employed PFA fixation as a one-step treatment, to reduce background SHG signals. The cilia in the hippocampal neuron greatly impact all biological functions of the brain, including spatial cognitive capacity and learning (25, 26). Figure 1a presents SHG-CARS images of the dentate gyrus (DG) region, where the CARS image was reconstructed from the Raman intensities at 2850 cm^-1^ that primarily correspond to CH stretching vibrations of lipids (see the Methods section for the detailed protocol). In the SHG image (Fig. 1a), only few-micrometer spots were sparsely detected without widespread SHG signals originating from axons, unlike the reported SHG studies on unfixed neurons (24). In the CARS image, the morphological features of the hippocampus were well contrasted, with the dark regions (shown in black) corresponding to the lipid-poor somatic layer and the bright regions (shown as white or cyan in the images) corresponding to the myelin sheath wrapping around axons; this was consistent with the results obtained by Evans et al. (27). As the merged (SHG (pink) and CARS (cyan)) image shows, the SHG signals were detected in the regions where the CARS signals were absent (the cell body regions); therefore, we conclude that the origin of the SHG signals are not axons. This background reduction of the axonal SHG signal is likely due primarily to the conformational change of tubulin proteins by PFA fixation (28), which is crucial for the successful visualization of the spot-like SHG signals. Building upon our previous spot-like SHG imaging study on the retina and COS7 cells (22), the ciliary rootlet emerges as a potential origin of the sparse SHG spots. Therefore, to test this hypothesis, we performed SHG/fluorescence imaging of a hippocampal tissue immunostained with anti-Rootletin, a marker of ciliary rootlets. We identified high colocalization of SHG spots and fluorescence spots (Fig. S1a), supporting our hypothesis that the SHG originates from ciliary rootlets. This is consistent with the spatial profile of the merged image, where only one SHG spot exists in each cell in the hippocampal somatic layer. Based on these experimental observations, we conclude that the SHG signals in Fig. 1a are assignable to the ciliary rootlet.

**Figure 1.**
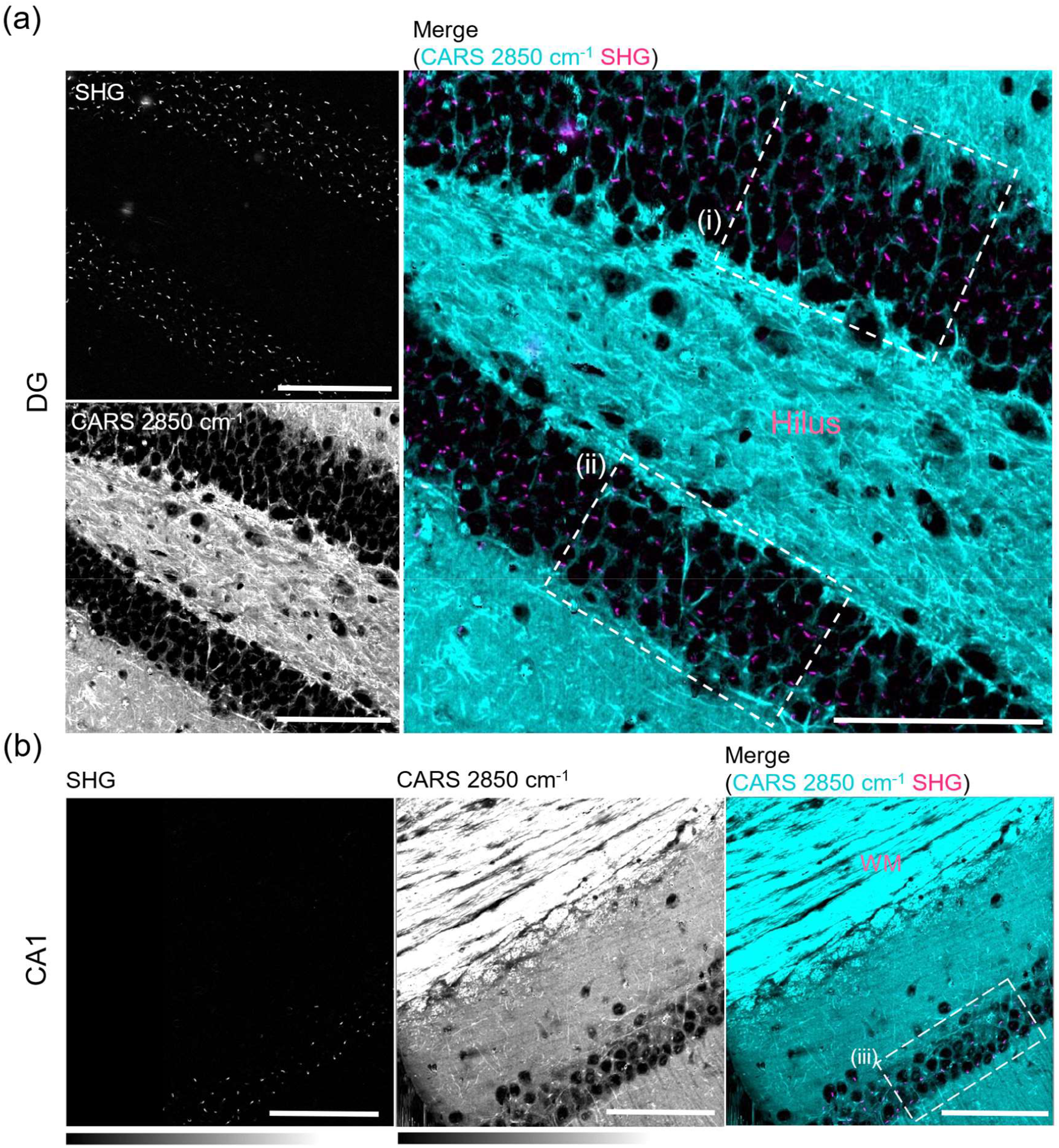
Label-free imaging of ciliary rootlets by SHG in the hippocampus. (a, b) SHG and CARS (2850 cm^−1^, CH_2_ symmetric stretching mode) and merged image in (a) Dentate gyrus and (b) CA1. The SHG spot is visible in each cell at the cell layer (dark regions). SHG and CARS visualize ciliary rootlets and myelin sheath, respectively. The image and step size used to acquire SHG and CARS images were 300 × 300 µm^2^ (601 × 601 pixels) and 0.5 µm/pixel, respectively (Scale bar, 100 μm). DG: dentate gyrus. WM: white matter.

We also conducted SHG-CARS imaging spanning the white matter (WM) and CA1 region for consistency in methodology (Figure 1b). The WM predominantly comprises axonal bundles, while the SHG signal derived from axons is typically abundant in this region. The spot-like SHGs were also observed in the somatic layer of the CA1 region (Fig. 1b). Similarly in this region, SHG/fluorescence imaging (Fig. S1b) confirmed that SHG originated from ciliary rootlet. In addition, the bright areas in the CARS image distinctly revealed the axonal bundles within the WM. However, no SHG signals were detected, which is consistent with the attenuation of the tubulin SHG signal by PFA fixation. This underscores the reproducibility of our methodology in the WM and CA1 regions.

### Ciliary rootlets are localized at the base of apical dendrites

We inadvertently discovered that the ciliary rootlets in the cell body are oriented in the same direction. Based on the large-area SHG-CARS images of the hippocampus, we statistically elaborated the intracellular positions of ciliary rootlets. The rootlet orientation in each cell was defined by the angle between the axis along the axons and the line connecting the centers of inertia of the cell and rootlet (Fig. 2a). To quantify the ciliary rootlet orientation of each cell, SHG and CARS images in the DG (Figs. 2b, 2c) and CA1 (Fig. 2d) regions were analyzed using image segmentation algorithms. The position of each rootlet was determined by the center of inertia of each bright SHG spot, whereas that of the cell was determined by each dark circle in the CARS images. In the cell clusters within the DG and CA1 regions, the ciliary rootlets were oriented opposite to the direction of axon outgrowth (Figs. 2e-2g). In both regions, the orientation distributions were maxima at 180°, the angle where the dendrites elongate with finite variances (Figs. 2e-2g). This *in vivo* result is similar to previous *in vitro* observations in cultured cells that showed that centrosomes are localized at the root of neuronal apical dendrites (29). Similarly, our results show that the ciliary rootlet extends from the basal body derived from the centrosome (17). Moreover, we found that even for healthy animals, certain cells showed irregular orientations at ∼0°, illustrating the importance of analyzing many cells to comprehensively understand the developmental processes of the brain and of using imaging data for ciliopathy diagnoses.

**Figure 2.**
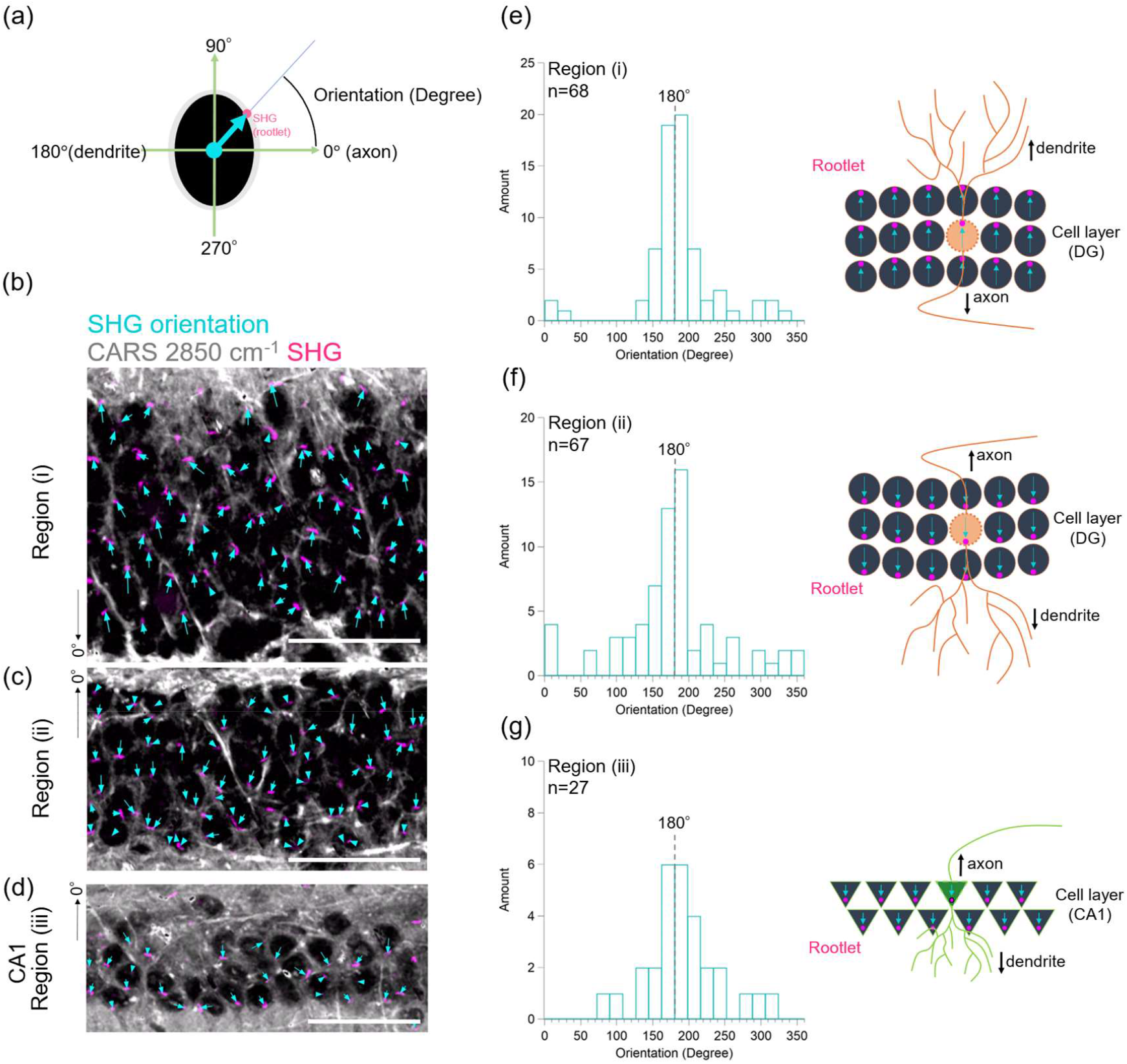
Orientation analysis of ciliary rootlets in the neuronal layer of DG and CA1. (a) Schematic diagram of the definition of orientation. Black: cell body (CARS at 2850 cm^−1^), pink: ciliary rootlet (SHG), cyan arrow: direction from the center of the cell body to ciliary rootlet. The ciliary rootlet orientation in each cell was defined by the angle between the axis along the axons and the line connecting the centers of inertia of the cell and rootlet (Defined all morphological axonal extensions as 0°). (b–d) Localization orientation analysis of SHG spots for each cell body in the tissue in Fig. 1(I–iii). (Scale bar, 100 μm). (e–g) Cell count distribution according to the angle of ciliary rootlets for each respective region (i–iii). Schematics show axonal and dendritic morphological extensions according to region. Ciliary rootlets are localized at the base of the dendrite in hippocampal neurons.

### Ciliary rootlets appear prior to the primary cilia extension

To determine the developmental stage of neurons at which ciliary rootlets first appear, primary cultured neurons at different stages isolated from the hippocampus were examined. Neurons are generated from the neuronal stem cells adjacent to the hilus in the hippocampus and gradually move toward the outer layer (30, 31). Therefore, the developmental stages differ by layer. The neurons in the layer adjacent to the hilus are predominantly immature, lacking cilia for the first five days post-differentiation (32). Interestingly, we observed Rootletin-originated SHG spots in the immature cell layer adjacent to the hilus. To investigate the behavior of ciliary rootlets in the early cell stage before the emergence of cilia, we performed SHG imaging of primary neurons cultured for 1 and 14 days. Moreover, we conducted immunostaining to compare the positioning with centrosomes, using anti-Pericentrin as a marker of centrosomes (Fig. 3). The comparison shows us that Rootletin proteins are localized adjacent to centrioles in immature neurons without cilia at div 1, meaning that Rootletin SHG signals are a precursor of cilia development. As the intracellular positions of the cilia and basal body are important markers of ciliopathies (33), these results suggest that SHG imaging is a promising tool for label-free, early diagnosis of ciliopathies.

**Figure 3.**
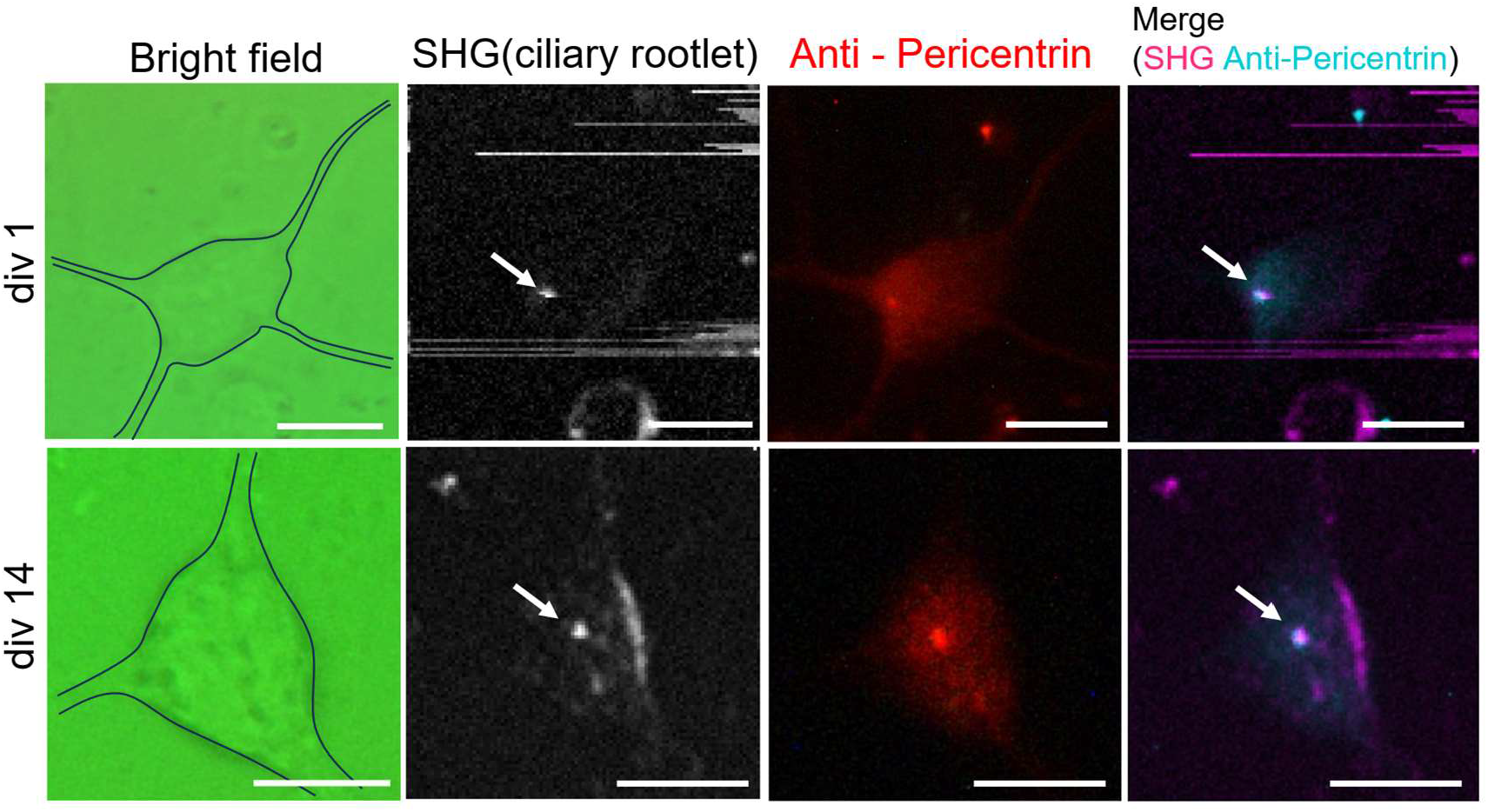
Comparison of ciliary rootlets in the developmental stages and immunostaining with anti-Pericentrin from the fixed primary culture neurons in the hippocampus. The image and step size used to acquire SHG were 30 × 30 µm^2^ (121 × 121 pixels) and 0.25 µm/pixel at div 1, and 25 × 25 µm^2^ (101 × 101 pixels) and 0.25µm/pixel at div 14, respectively(Scale bar, 10 μm). Ciliary rootlets (white arrow) co-localize with centrosome and have existed since div 1. div: day of *in vitro*.

### Molecular structure of ciliary rootlets is similar to myosin

To further elucidate the molecular structural characterization of rootlets, we applied our SHG-CARS imaging technique to ependymal cells with high-density rootlets in the cerebral ventricle (Fig. 4). Ependymal cells drive cerebrospinal fluid circulation by multiple cilia formed by centriole replications independent of cell division (Fig. 4a). Considerably stronger SHG signals were observed than in the hippocampus (Fig. 4b), which is consistent with Rootletin abundance. Moreover, SHG/fluorescence imaging confirms the origin of the SHG signal in this region as Rootletin (Fig. S3). Moreover, CARS images around the cerebral ventricle at 1660, 1546, 1401, and 1043 cm^-1^, which are assignable to amide I, tryptophan, COO-antisymmetric stretching, and proline, respectively (Fig. 4b). For these four peaks, relatively intense CARS signals were detected in the surface region, where strong SHG signals were detected. This colocalization of the SHG- and CARS-active regions facilitates further investigation of the molecular structure of Rootletin proteins in the SHG-active region. Additionally, CARS spectra exhibited Raman bands at 1546, 1401, 1307, 1255, 1201, and 1043 cm^-1^ in the SHG-active region (Fig. 4c-i) not throughout the entire tissue (Fig. 4c-ii). These bands are also present in myosin’s Raman spectrum (34), implying a common lineage and molecular structural similarity between myosin and Rootletin. In fact, the evolutionary strains of the two are similar (17). While there are reports of non-muscle myosin in ependymal cells (35), similar reports for SHG have not been made. Given that the SHG images mirrored the Rootletin immunostaining images (Fig. S2a), we suggest that the CARS spectrum averaged over the bright SHG areas is derived from Rootletin.

**Figure 4.**
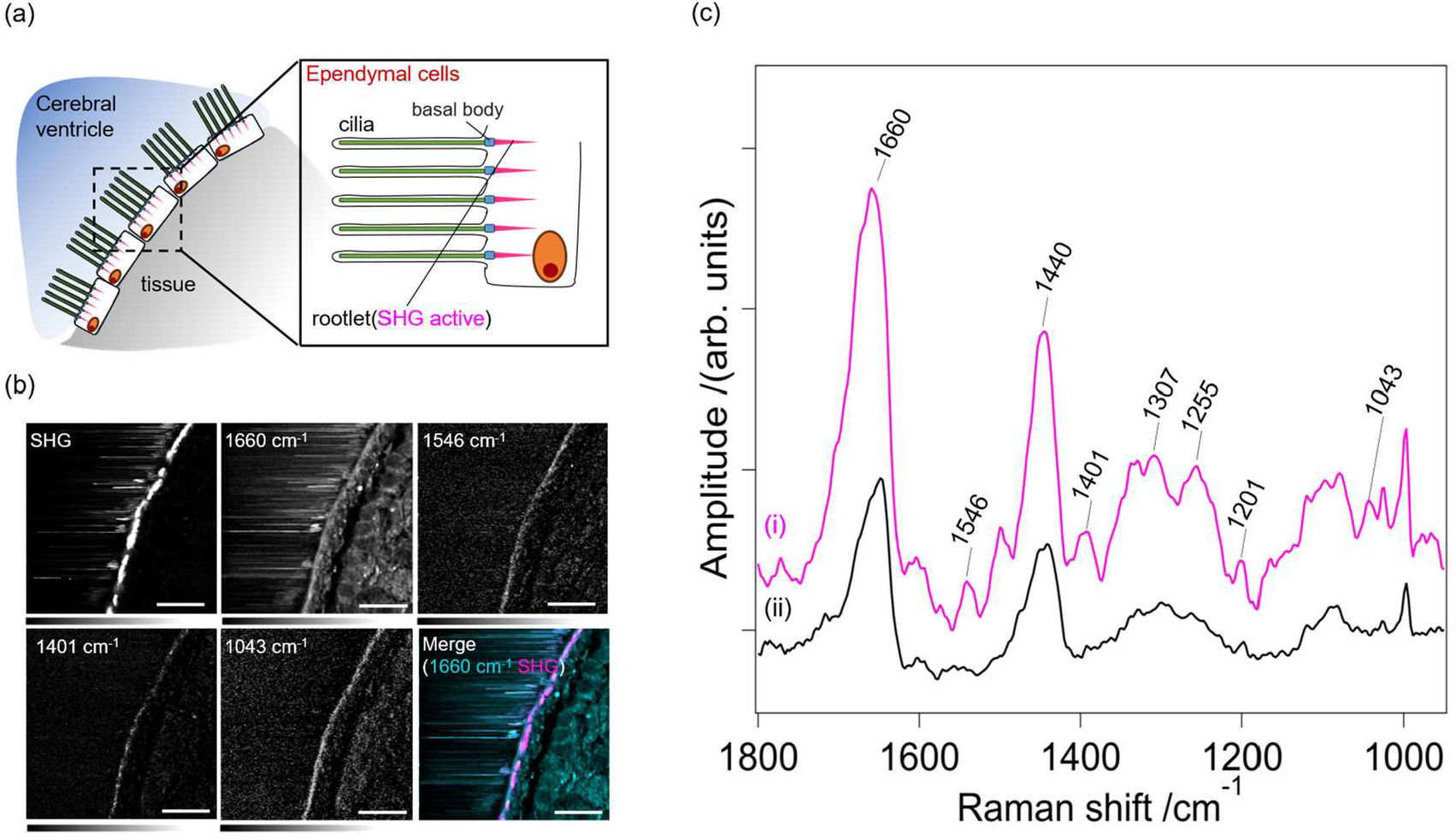
SHG and CARS imaging of ependymal cells around the cerebral ventricle. (a) Schematic of the area around the cerebral ventricle. The ependymal cells are dense in the cerebral ventricles and elongate the multiple cilia and corresponding ciliary rootlets. (b) SHG, CARS at 1660, 1546, 1401, and 1043 cm^−1^, and merged image of surrounding cerebral ventricle in mouse brain. The image and step size used to acquire SHG and CARS images were 100 × 100 µm^2^ (201 × 201 pixels) and 0.5 µm/pixel, respectively. (scale bar: 25 µm). (c) CARS spectra of SH active region (i) and average spectra of tissue (ii).

## Discussion

Our label-free imaging technique is expected to serve as a valuable new tool for exploring ciliary rootlets in various biological settings. Rootletin maintains its structural integrity even under PFA fixation, similar to myosin, as evidenced by the absence of SHG intensity attenuation, offering the potential for broad application across biological samples, including human tissue diagnostics. Furthermore, our imaging platform shows promise in elucidating the localization of ciliary rootlets and enhancing our understanding of brain development. Visualization of Rootletin fiber is also useful for label-free imaging of centrosomes. Variations in centrosome positioning correspond to the direction of cell motility and the position of cilia, promoting axon growth and determining the direction of axons in mammalian neurons (33, 36). Meanwhile, centrosome mispositioning has been linked to ciliopathies (33). Our method allows for the simultaneous observation of centrosome and ciliary rootlet orientation heterogeneity across multiple cell types, providing insights regarding the diverse cellular interactions underlying disease development. We also observed the presence of ciliary rootlets *before* ciliogenesis. However, the role(s) of ciliary rootlets prior to ciliogenesis remains elusive, warranting further examination to better characterize their contributions to neuronal differentiation and diseases.

SHG imaging also has the potential to facilitate dynamic observations of ciliary rootlets *in vivo* via epi-detection. SHG has facilitated the imaging of collagen and myosin *in vivo*, as well as the detection of backward signals, which are essentially weaker than forward signals, and improve with heterodyne-enhanced signal amplification (37). Another issue in extending the technique to epi-detection is the transparency of the excitation light. Currently, *in vivo*, two-photon fluorescence imaging in the hippocampus has been successful due to the improved bio-transparency by appropriate wavelength selection and algorithms (38-40). The SHG setup is nearly equivalent to two-photon imaging and should be equally feasible for *in vivo* extension. In this case, the unfixed distinction between tubulin and rootlets is essential and is likely distinguishable by the polar angle of the harmonophore (θ_e_). We previously found that the polar angle of the Rootletin harmonophore (θ_e_ = 55.5 ± 3.6°) differs from that of collagen (θ_e_ = 62° to 69°) (22). Furthermore, the distribution of the effective orientation of the SHG active source in cultured primary cortical neurons is reportedly centered at θ_e_ = 33.96° (41). This suggests that the polarization dependence of axons and Rootletin may differ, allowing for the distinction by the polarization direction of the excitation laser. Furthermore, CARS (corresponding to lipids in the CH_2_ stretching mode) and THG signals are active in the myelin sheath (27, 42). However, they are active in axons and inactive in ciliary rootlets, potentially allowing for selective distinction of ciliary rootlets. Therefore, extending our method to *in vivo* imaging allows for more natural, tissue-level observation of ciliary rootlet dynamics and can potentially facilitate the comprehensive elucidation of diseases, including ciliopathies.

## Materials and methods

### SHG and CARS imaging system

A custom-made dual-fiber, output-synchronized light source (OPERA HP; Leukos, Limoges, France) was used. One of the fiber outputs provided laser pulses with a 1064 nm wavelength, 50 ps pulse width, and 1 MHz repetition rate. They were used as the pump beam (ω_1_) for the CARS process. The other fiber outputs provided supercontinuum (SC) radiation ranging from visible to near-infrared (NIR) with a broad wavelength of 1100–1800 nm. The SC radiation was spectrally filtered using a long-pass filter (IR80, Kenko-optics, Tokyo, Japan) and applied as the Stokes beam (ω_2_). The pump and Stokes laser beams were superimposed using a notch filter (NF03-532/1064E-25; Semrock, Rochester, NY, USA) and guided to a modified inverted microscope. Two laser pulses were tightly focused on the sample using a microscope objective (CFI Plan Apo 60x NA 1.27, water immersion, Nikon). Samples were placed on a piezoelectric stage (Nano-LP300; Mad City Labs, Madison, WI, USA) for position selection. The full scanning range of the XYZ piezo stage was 300 μm^3^. In addition, CARS and SHG signals were collected using a second objective lens (Plan S Fluor 40 × NA0.6, Nikon). The CARS (2ω_1_ - ω_2_) and SHG (2ω_1_) signals were spectrally separated by a dichroic mirror. The SHG signals were detected using a spectrometer (SpectraPro300i; Princeton Instruments Inc., Trenton, NJ, USA) equipped with a CCD camera (PIXIS 100 B; Princeton Instruments). The CARS signal was detected using a spectrometer (LS785, Princeton Instruments) equipped with a CCD camera (BLAZE 100HR, Princeton Instruments). The CARS (Im[c^(3)^]) spectrum, equivalent to the spontaneous Raman spectrum, was retrieved from the raw CARS spectrum using MEM (43).The spectral coverage and spectral resolution of the CARS signal were approximately 3500 and 8 cm^−1^, respectively. The exposure time for each pixel was 100 ms. The laser powers were approximately 200 mW and 170 mW for the pump and Stokes, respectively.

### Multi-modal (SHG and TPEF) imaging by 1064 nm excitation for primary culture neuron and mouse brain slice

A microscope equipped with a pulsed laser (OPERA HP2; Leukos, Limoges, France) served as the light source. The OPERA HP2 had a laser output wavelength of 1064 nm, pulse width of 50 ps, and repetition rate of 5 MHz. This was used as the excitation beam for the SHG process. The exposure time for each pixel was 200 ms. The laser power was approximately 200 mW for excitation in the sample plane.

In addition, the same light source was used to observe two-photon excitation fluorescent (TPEF) for the molecular identification of SHGs in the cerebral ventricle. Figure S3 shows the spectral profile of the immunostained cerebral ventricle at the position of the SHG filament with 1064 nm laser excitation. A sharp and intense band at 532 nm and a broad and weak band at approximately 570 nm were observed, corresponding to SHG and TPEF, respectively. The bands at approximately 570 nm were assigned to TPEF owing to Alexa Fluor 546, which stained the tissue. Notably, the TPEF signal was detectable in the cerebral ventricle but not in cultured cells or the cell layer of the hippocampus. This is because multiple cilia densely populate the ependymal cells at the boundary of the cerebral ventricle.

The TPEF signals of Alexa Fluor 546 were difficult to detect in cultured neurons or the cell layer of the hippocampus. To observe fluorescently stained cells within neurons for comparison of fluorescence images and SHG, the SHG system was increased with a mercury lamp (Intensilight C-HGFIE, Nikon, Tokyo, Japan), appropriate excitation and emission cubes (TRITC, Nikon, Tokyo, Japan), and a dedicated CCD camera (Thorlabs, 1500M-GE).

### Preparation of mouse brain samples

Coronal brain sections from the wild-type mice (C57BL/6 N) were used as samples for multimodal imaging. Male mice at p97 were deeply anesthetized with 5% isoflurane and somnopentyl isoflurane (32.4 mg/kg). Thereafter, the mouse was transcardially perfused with 0.1 M phosphate-buffered saline (PBS) with heparin for 1 min, followed by transcardial perfusion with 4% paraformaldehyde for 10 min. The brain was removed and immersed in the same post-fixation buffer at 4 °C for 24 h. The removed brain was washed thrice with PBS (5 min each, shaking in a shaker) and subsequently sectioned into 100 µm thick sections using a vibratome (Leica, VT1000S). The resulting brain sections were subjected to label-free CARS and SHG imaging.

### Cell culture

Dissociated rat hippocampal neurons, together with culture media, were purchased from Lonza Bioscience (Walkersville, MD, USA) and used according to the manufacturer’s instructions. Briefly, a frozen stock of rat hippocampal neurons was thawed and diluted in 16 mL of culture medium, which was then plated into eight 35 mm dishes, each containing four collagen-coated coverslips (AGC Techno Glass, Shizuoka, Japan) and additionally coated with poly-L-lysine. After 4 h of incubation in a 37 °C 5% CO_2_ incubator, the entire medium was changed to fresh media. Neurons were maintained in a 37 °C 5% CO_2_ moist incubator until use, with half of the medium changed twice weekly.

### Antibodies and Immunostaining

First, 100 μm-thick fixed coronal brain slices were incubated with permeabilization and blocking solution containing 10% normal goat serum and 0.5% triton X-100 in PBS for 30 min at room temperature (RT). Brain slices were then incubated with an anti-Rootletin rabbit antibody (Millipore ABN1714, Burlington, MA, USA) diluted in antibody dilution solution containing 2% normal goat serum and 0.1% triton X-100 in PBS overnight at RT. Brain slices were washed three times in PBS and incubated with a secondary antibody (Alexa Fluor 546 conjugated anti-rabbit goat antibody, Thermo Fisher Scientific, Waltham, MA, USA) in antibody dilution solution for 1 h at RT, followed by washing in PBS thrice. Immunostaining of primary culture neurons was performed in the same manner using an anti-Pericentrin antibody (ab4448, Abcam).

### Quantification of rootlet orientation in the brain

The analysis was performed using custom programs in ImageJ and Fiji. This algorithm defines regions of interest (ROIs) for cell bodies and the ciliary rootlet (SHG) indicated by CARS, determining the center pixel of each ROI. Subsequently, only discernible cells were arbitrarily matched between the cell body and rootlet ROI, connecting their centers with line segment ROIs (from the center of the cell body ROI to the center of the ROI, indicating their orientation). The angles of all line segment ROIs were quantified using angles defined within ImageJ; the axon extension direction was defined as 0, with 360° maximum counterclockwise in each region. Cyan arrows indicate the ROI of the line segment and its direction (i.e., localization direction).

## Acknowledgments

The authors thank Dr. S. Miyazaki, Ms. M. Masaki, Ms. S. Tokunaga, and Mr. R. Sakamoto for their helpful discussions. This study was financially supported by the Japan Society for the Promotion of Science (JSPS) through a Grant-in-Aid for Scientific Research (A) (KAKENHI, grant number 21H04961), Japan Science, the Technology Agency (JST) through a Mirai Program grant (JPMJMI22G5) and AMED under Grant Number JP21zf0127005. It also benefited from a Japan–France bilateral project awarded to H.K. and the support of the French government in the form of a grant managed by the National Research Agency under the Investments for the Future program (ANR-10-LABX-0074 Sigma-LIM) awarded to P. L. The authors are grateful to J. Ukon of Ukon Craft Science, Ltd. for establishing the collaboration between the Japanese and French laboratories.

## Notes

### Competing Interest Statement

The authors have declared no competing interest.

