## Supporting Information for "Label-free visualization of ciliary rootlets in mouse brain"

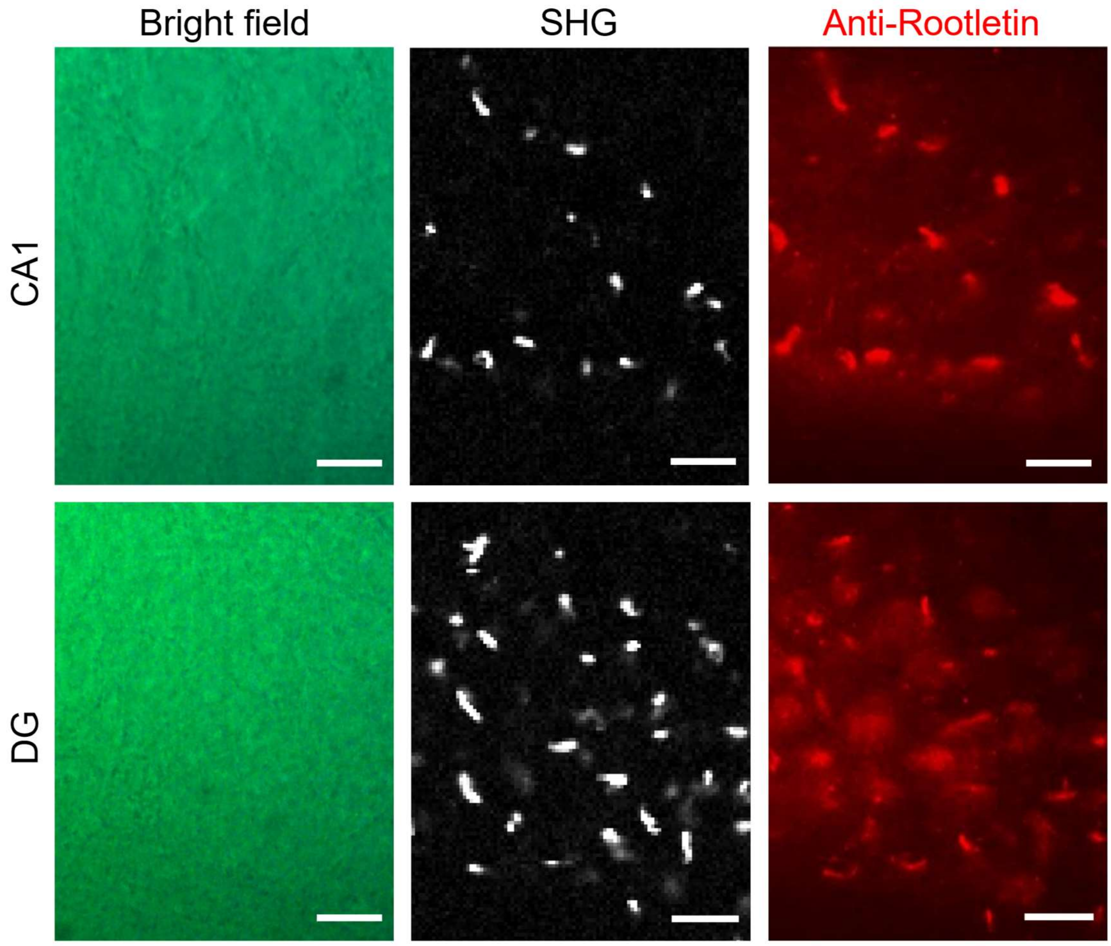


**Supplementally Figure 1.** Colocalization analyses of Second Harmonic Generation (SHG) and immunostaining with anti-Rootletin in the CA1 and DG regions of the hippocampus in fixed mouse brain. The image and step size used to acquire SHG were 50 × 60 $mm^2$ (100 × 120 pixels) and 0.5 μm/pixel, respectively(Scale bar, 10 μm). SHG and Rootletin were co-localised.

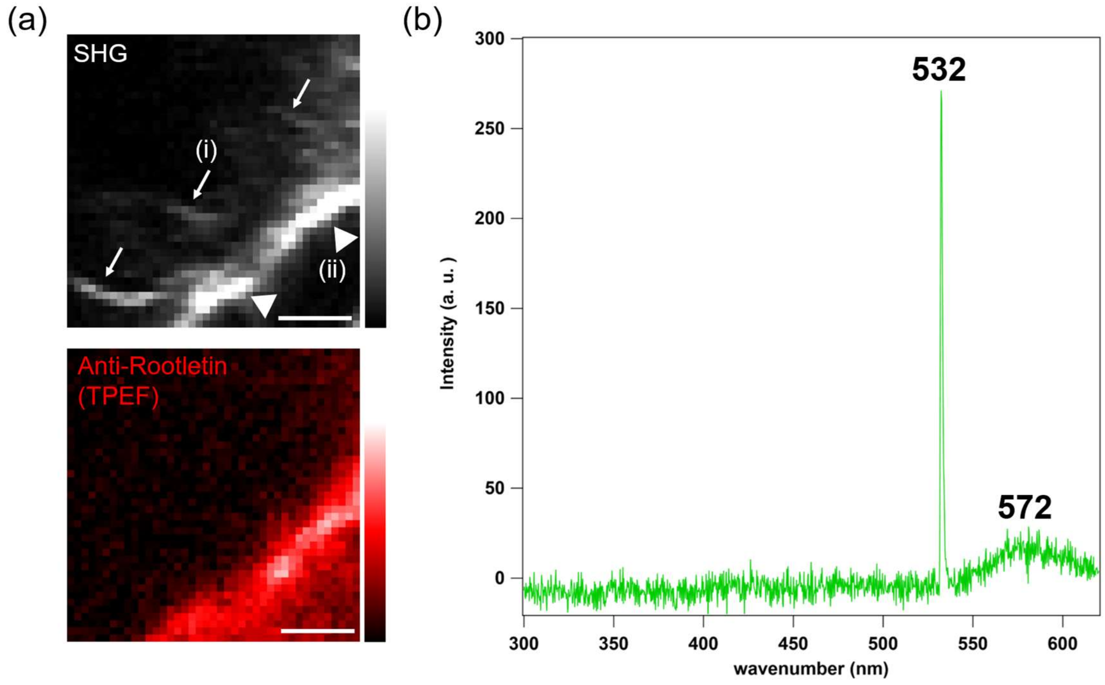


**Supplementally Figure 2.** SHG and TPEF from around cerebral ventricle. Spectral profile of immunostained mouse brain at the position of the Rootletin. The sharp and intense band at 532 nm and the broad and weak band around 572 nm correspond to SHG and TPEF, respectively. The bands around 572 nm were assigned as Alexa Fluor 546. (i) and (ii) are SHG signals derived from cilia and ciliary rootlets, respectively.
